# AF10 (MLLT10) prevents somatic cell reprogramming through regulation of DOT1L-mediated H3K79 methylation

**DOI:** 10.1101/2020.12.22.423908

**Authors:** Deniz Uğurlu-Çimen, Deniz Odluyurt, Kenan Sevinç, Nazlı Ezgi Özkan-Küçük, Burcu Özçimen, Deniz Demirtaş, Eray Enüstün, Can Aztekin, Martin Philpott, Udo Oppermann, Nurhan Özlü, Tamer T. Önder

## Abstract

The histone H3 lysine 79 (H3K79) methyltransferase DOT1L is a key chromatin-based barrier to somatic cell reprogramming. However, the mechanisms by which DOT1L safeguards cell identity and somatic-specific transcriptional programs remain unknown. Here, we employed a proteomic approach using proximity-based labeling to identify DOT1L-interacting proteins and investigated their effects on reprogramming. Among DOT1L interactors, suppression of AF10 (MLLT10) via RNA interference or CRISPR/Cas9, significantly increases reprogramming efficiency. In somatic cells and induced pluripotent stem cells (iPSCs) higher order H3K79 methylation is dependent on AF10 expression. In AF10 knockout cells, re-expression wildtype AF10, but not a mutant defective in DOT1L binding, rescues overall H3K79 methylation and reduces reprogramming efficiency. Transcriptomic analyses during reprogramming show that AF10 suppression results in downregulation of fibroblast-specific genes and accelerates the activation of pluripotency-associated genes. Our findings establish AF10 as a novel barrier to reprogramming by regulating H3K79 methylation and thereby sheds light on the mechanism by which cell identity is maintained in somatic cells.

## Introduction

The low efficiency of transcription factor-based reprogramming points to the presence of multiple rate-limiting steps or barriers to cell fate changes (Takahashi and Yamanaka, 2006). We have previously identified the histone H3 Lysine 79 (H3K79) methyltransferase DOT1L as one of the key barriers to reprogramming of somatic cells to pluripotency (Onder et al., 2012). DOTL1 inhibition can functionally replace KLf4 and c-MYC (Onder et al., 2012), increase reprogramming efficiency in a wide range of systems (Ebrahimi et al., 2019; Ichida et al., 2014; Jackson et al., 2016; Tran et al., 2019), facilitate the generation of chemically induced pluripotent stem cells (ciPSCs) from mouse somatic cells (Zhao et al., 2015) and result in a permissive epigenome state which enables reprogramming by alternative transcription factors (Kim et al., 2020). DOT1L is recruited to RNAPII-associated transcription-elongation machinery through a number of interacting proteins that include members of AEP (AF4 family/ENL family/P-TEFb), EAP (ENL-associated proteins), DotCom, and super-elongation protein complexes (Lin et al., 2010; Mohan et al., 2010; Mueller et al., 2009; Yokoyama et al., 2010). H3K79 methylation decorates actively transcribed gene bodies where it can act as an anti-silencing mark and prevent the recruitment of repressive chromatin modifiers (Chen et al., 2015a; Kouskouti and Talianidis, 2005; Steger et al., 2008; Stulemeijer et al., 2011). In the context of reprogramming, DOT1L activity serves to maintain the expression of somatic-specific genes and prevents mesenchymal-to-epithelial transition (MET), an important step in the process (Onder et al., 2012). However, the key interaction partners of DOT1L which play a role in safeguarding somatic cell identity remain unknown. In the present work, we addressed this question using a combination of proteomics and loss of function approaches and identified AF10 as a key DOT1L-interacting protein in maintaining cell identity.

## Results

### Identification of proximal interactors of DOT1L via BioID

To identify interaction partners of DOT1L in somatic cells, we generated a fusion protein linking a promiscuous biotin ligase (BirA*) with DOT1L (**Figure 1A**)(Roux et al., 2012). We also generated a BirA*-fusion with a catalytically dead DOT1L mutant (G163R/S164C/G165R) incapable of H3K79 methylation to assess if putative interactors could be dependent on catalytic activity of DOT1L (**Figure 1A**) (Okada et al., 2005). To test the functionality of these fusion proteins, constructs were transfected into control and DOT1L knockout 293T cells generated via CRISPR/Cas9. In the DOT1L knockout background, H3K79 methylation was restored upon expression of wild-type, but not mutant DOT1L fusion protein, confirming that BirA-fusion does not interfere with catalytic activity (**Figure 1B**). Biotinylated proteins were analyzed in LC-MS/MS. Mass spectrometry analysis resulted in detection of DOT1L with the highest PSM (peptide spectrum matches) values (1% false discovery rate (FDR)) and high sequence coverage (30%) in fusion protein-expressing samples; whereas none was detected in control samples as expected. In wt-DOT1L fusion expressing samples, 11 proteins were identified (**Figure1C**). Among these were a number of previously characterized interactors such as AF10, AF17, ENL as well six novel putative proximal-interactor proteins (TPR, KAISO, NUMA1, MRE11, NONO, SIN3B). In contrast, 96 proximal interactors were detected in mut-DOT1L expressing cells (**Figure 1C**). This larger number of biotinylated proteins in mut-DOT1L samples may be due to a defect in chromatin localization of the mutant protein, a notion that needs further investigation. We next asked whether any of the putative interactors of wt-DOT1L have an effect on the reprogramming of human fibroblast to iPSCs. In a loss of function approach, we knocked-down individual candidate genes by two independent shRNAs. The majority of shRNAs achieved at least 50% knock-down of their respective target gene (**Figure S1A**). Reprogramming was initiated after shRNA transduction and the resulting iPSC colonies were identified via Tra-1-60 expression, a well-established marker of fully reprogrammed cells (Chan et al., 2009). We observed that knock-down of *AF10* and *NONO* significantly increased the number of iPSC colonies, resulting in 1.5 to 2-fold greater reprogramming efficiency compared to control shRNA expression (**Figure 1D**). On the other hand, knock-down of *MRE11* and *TPR* decreased reprogramming significantly (**Figure 1D)**.

**Figure 1:**
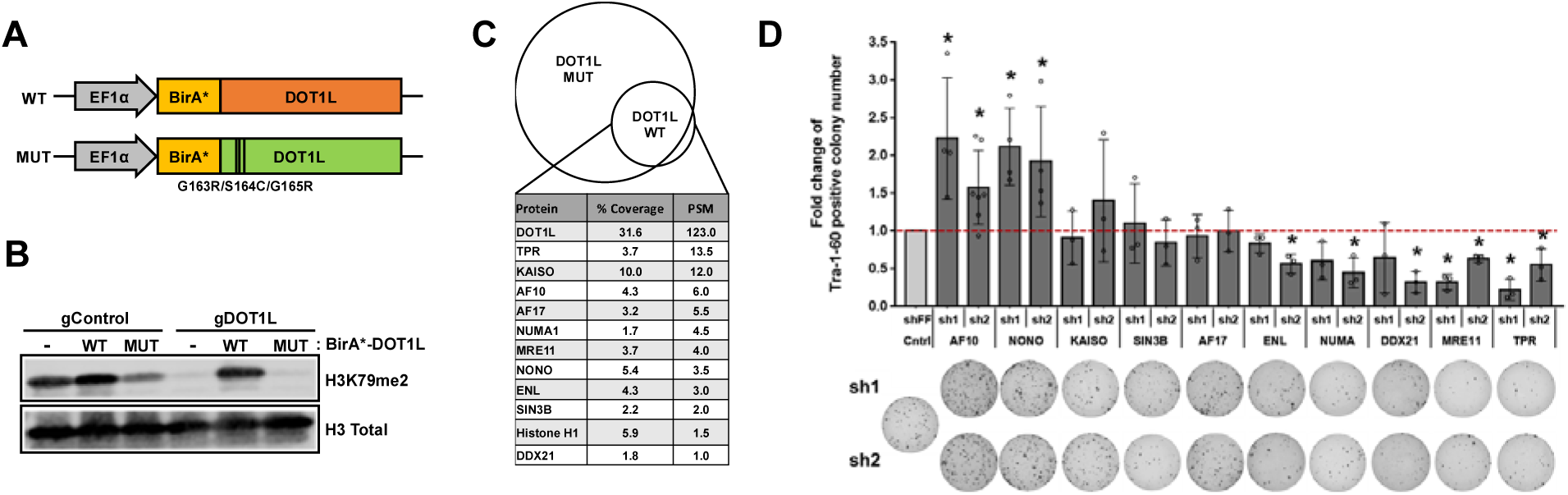
Identification of proximal interactors of DOT1L via BioID. **(A)** Schematic of BirA*-DOT1L fusion protein expressing vector constructs. **(B)** H3K79 di-methylation (H3K79me2) levels in control (gControl) and DOT1L knock-out (gDOT1L) cells expressing either WT or MUT BirA*-DOT1L fusion constructs. Total histone H3 was used as a loading control. **(C)** Proximal-protein interactions of DOT1L as revealed via proteomic analyses of biotinylated proteins. Venn diagram represents the number of biotinylated proteins after mass spectrometry analysis. Proteins were ordered according to their average coverage and PSM (peptide spectrum matches) values in BirA*-DOT1L wt expressing cells. **(D)** Bar graph represents fold change in reprogramming efficiency upon shRNA-mediated gene silencing. Tra-1-60 positive colony numbers of each experiment were normalized to shControl sample. Average of fold changes from independent experiments are indicated (circles). Representative Tra-1-60 stained well images for each shRNA-infected sample are displayed under the bar graph. Error bars represent SEM. *, P < 0.05.

### AF10 suppression enhances reprogramming

We were intrigued by the increased reprogramming efficiency upon *AF10* and *NONO* knock-down and followed up on these two candidate genes. We next asked if these two proteins play a role in regulating cellular H3K79 methylation levels. Knock-down of *NONO* did not change total H3K79me2 levels (**Figure S1B**). Considering that Nono has been shown to limit self-renewal of mESCs by regulating bivalent gene expression, reprogramming enhancement upon *NONO* knockdown may occur independent of H3K79 methylation (Ma et al., 2016). In contrast, AF10 inhibition via shRNAs significantly decreased H3K79 methylation (**Figure S1B**). To further confirm the role of AF10 in reprogramming, we pursued an independent strategy to inhibit AF10 using two independent guide RNAs (gRNAs) targeting splice site exon 2 or exon 3 of AF10 (*MLLT10*) (Chen et al., 2015b) (**Figure 2A**). CRISPR-targeted sites were verified via T7 endonuclease assay and sgAF10-expressing fibroblasts had lower *AF10* mRNA levels compared to sgControl-expressing cells (**Figure 2A, S1C**). In addition, H3K79 methylation was decreased in both sgAF10 cell lines, albeit to a lesser degree than treatment with a small molecule inhibitor of DOT1L (iDOT1L, EPZ004777) (**Figure 2C**). sgAF10 expressing-fibroblasts generated up to 2-fold greater number of iPSC colonies compared to control cells (**Figure 2D**). We next evaluated if iPSCs derived via AF10 suppression were *bona fide* pluripotent cells. AF10 and H3K79me2 levels were significantly reduced in all sgAF10-derived iPSC clones tested (**Figure 2E**). sgAF10 iPSC colonies were positive for OCT4, SSEA4 and NANOG at the protein level, and, upon injection into immunodeficient mice, readily formed teratomas containing cells originating from all three germ layers (**Figure 2F, G**). Overall, these experiments show that cells with AF10 inhibition can be fully reprogrammed into *bona fide* iPSCs.

**Figure 2:**
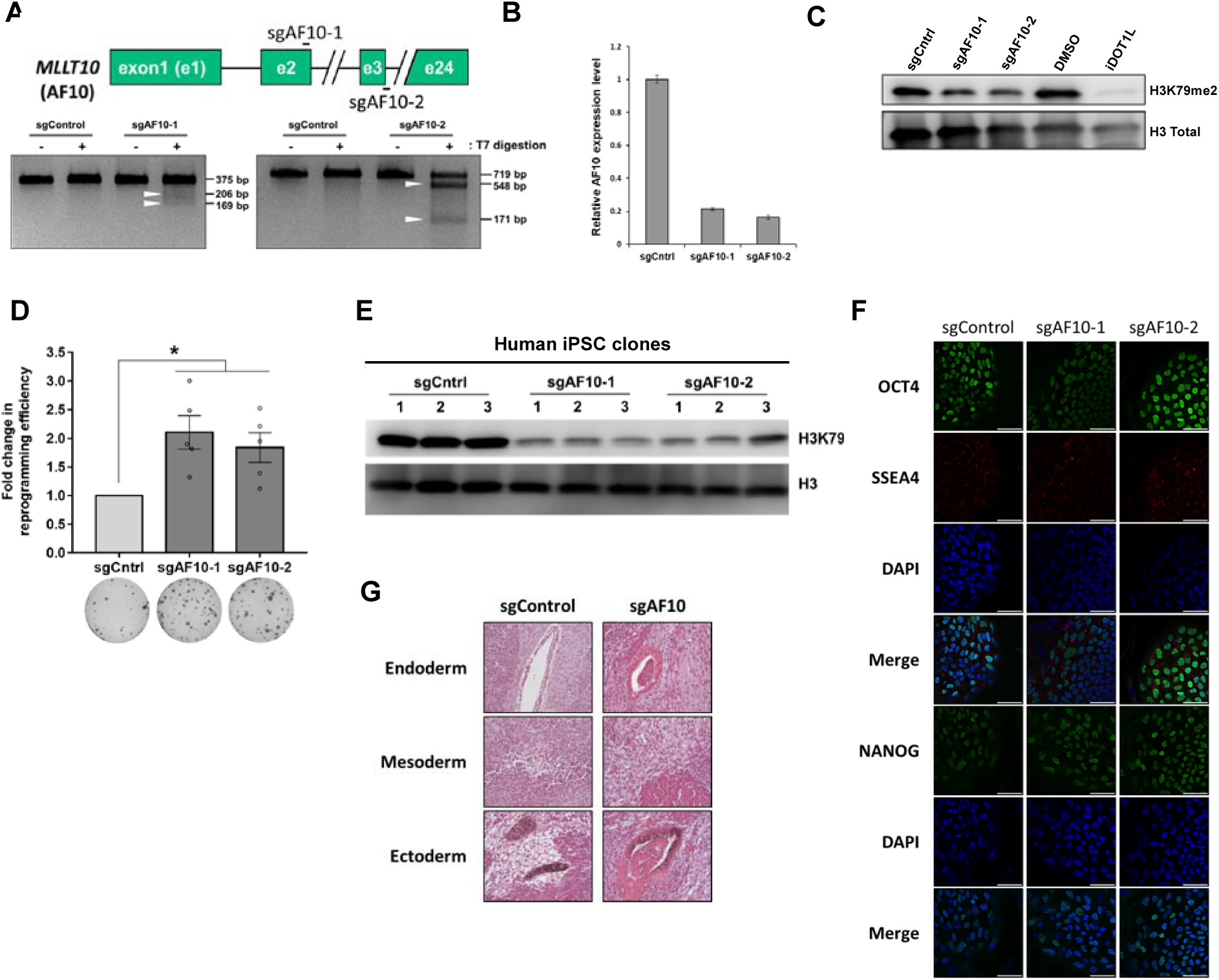
AF10 regulates H3K79 methylation and is a barrier to reprogramming. **(A)** Schematic for AF10 (MLLT10) gene indicating target sites for the AF10 sgRNAs. T7-endonuclease assay for sgAF10 target sites (bottom). Expected DNA fragments are indicated with white arrow heads. **(B)** AF10 mRNA levels in control and sgAF10 expressing cells as determined by qRT-PCR. β-actin was used as an internal control and expression level is normalized to sgControl expressing cells. **(C)** H3K79me2 uponn sgRNA-mediated AF10 knockout. iDOT1L (EPZ004777) was used as a positive control of H3K79me2 depletion. Fibroblasts were treated with DMSO or 3 μM iDOT1L for 10 days. sgAF10 infected dH1fs were selected with puromycin and cultured for 1 week. Total H3 levels are used as loading control. **(D)** Fold change in the number of Tra-1-60 positive colonies upon sgAF10 expression. P values were determined by one sample t-test; * P⍰< ⍰0.05. Bar graphs show the mean and error bars represent SEM in independent biological replicates (each circle). Representative Tra-1-60 stained wells are shown below the graph. *P* values were 0.009 for sgAF10-1 and 0.016 for sgAF10-2. **(E)** Immunoblot for H3K79me2 in individual control and sgAF10 iPSC lines. Total H3 levels were used as loading control. **(F)** OCT4, SSEA4 and NANOG immunofluorescence of iPSCs derived from control and sgAF10 expressing fibroblasts. DAPI was used to stain the nuclei. Scale bars represent 50 μm. **(G)** Hematoxylin and eosin stained sections of teratomas generated by iPSCs derived from control and AF10 knockout cells. Panels show glandular epithelium (endoderm, top), cartilage tissue (mesoderm, middle), and pigmented neural tissue (ectoderm, bottom). Representative images are from one of two independent teratomas.

We next asked whether the increased reprogramming phenotype upon AF10 knock-out could be rescued by re-expression of AF10. Wildtype *AF10* cDNA increased overall H3K79me2 levels in sgAF10-expressing cells, and importantly, decreased the reprogramming efficiency (**Figure 3A-C**). Thus, the increased reprogramming phenotype upon AF10 silencing could be rescued by overexpression of WT-AF10. Using the same approach, we next asked if a H3K27 binding-mutant of AF10 (L107A) and a DOT1L-binding domain deleted AF10 (octapeptide motif-leucine zipper deletion, OM-LZΔ) would behave similarly in reprogramming. The increased reprogramming phenotype was reverted by the L107A but not the OM-LZΔ mutant, indicating that AF10-DOT1L interaction, but not histone binding, is critical (**Figure 3C**). These results altogether show that AF10 constitutes a barrier to reprogramming to pluripotency and that its binding to DOT1L is important for this function.

**Figure 3:**
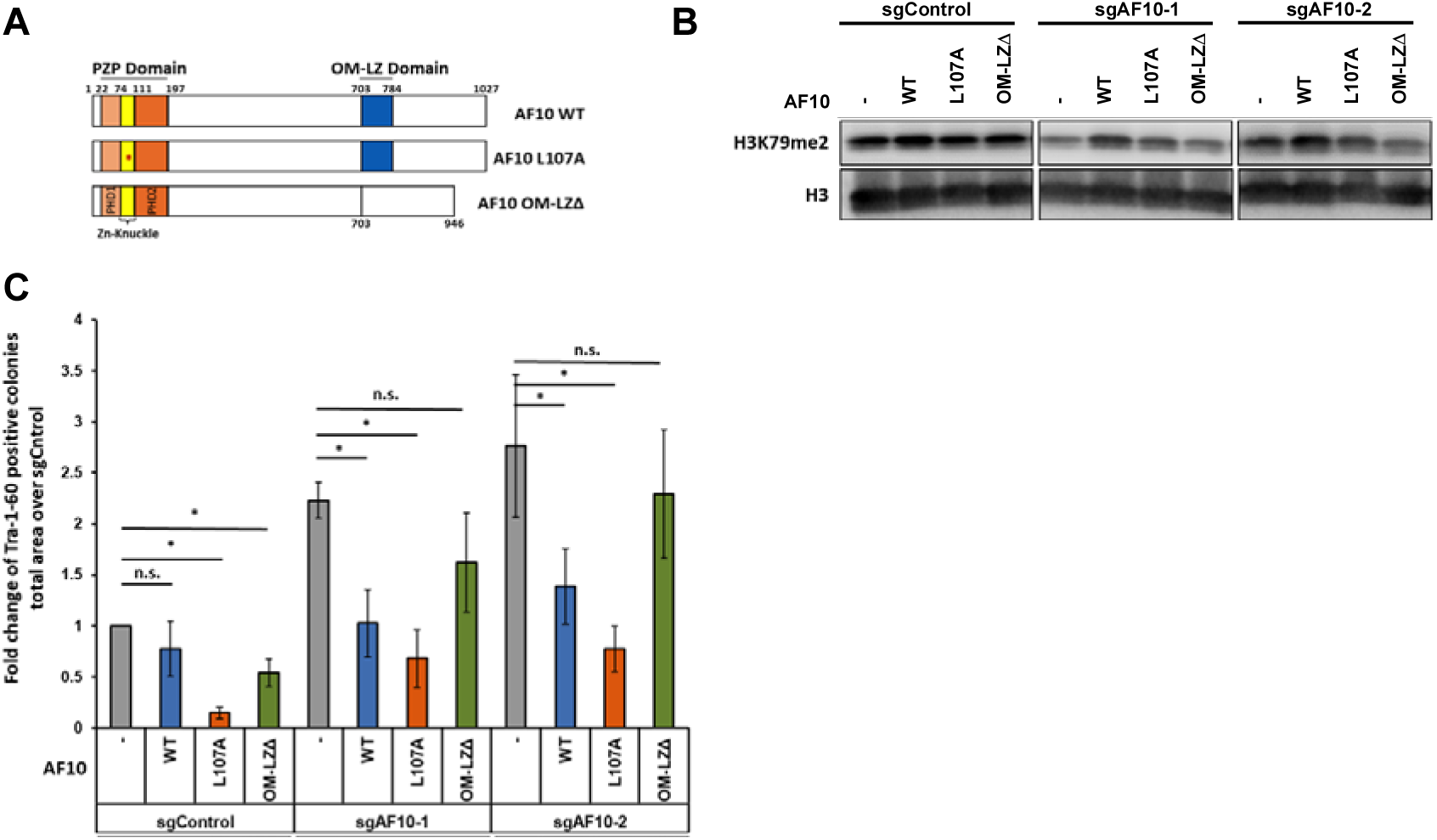
AF10 prevents reprograming through its interaction with DOT1L **(A)** Domain organization of wildtype and mutant AF10s used in (A). L107A mutation and OM-LZ deletion abolishes Histone H3 or DOT1L binding, respectively. **(B)** Immunoblots for H3K79me2 levels in control and AF10 knock-out cells expressing either wildtype or mutant AF10 cDNAs. Total H3 levels were used as loading controls. **(C)** Fold change in the number of Tra-1-60-positive colonies derived from control or AF10 knockout cells expressing wt, L107A or OM-LZ? mutant AF10 cDNAs. *P* values were determined by one sample t-test; * P⍰< ⍰0.05. Bar graph shows the mean and error bars represent SEM in 3 independent biological replicates.

### AF10 expression maintains somatic cell identity

To elucidate the mechanism by which AF10 suppression enhances iPSC generation, we investigated the transcriptional changes occurring upon sgAF10 expression. Since AF10 loss has a clear effect of H3K79me2 levels, we hypothesized that it will affect the transcriptional landscape of somatic cells. We performed an RNA-sequencing experiment in control and sgAF10-expressing cells early during reprogramming, on day 6 post-OSKM expression. A large number of genes were differentially expressed between control and sgAf10 expressing fibroblasts upon OSKM induction (749 genes upregulated; 735 genes downregulated). We specifically asked whether pluripotency-associated genes were upregulated upon AF10 inhibition. Gene-set enrichment analysis (GSEA) indicated that pluripotency genes were highly enriched in sgAF10 cells upon OSKM expression (**Figure 4A**). On the other hand, fibroblast-related genes were negatively enriched upon sgAF10 treatment, which suggested greater suppression of the somatic cell-specific gene expression program (**Figure 4A**). We next assessed the degree to which AF10 and DOT1L-induced transcriptional changes overlap during reprogramming. Based on published gene expression data of DOT1L inhibitor-treated cells, we generated gene sets comprised of genes negatively or positively regulated by DOT1L (Onder et al., 2012). GSEA of sgAF10 transcriptome data revealed that iDOT1L-downregulated genes were negatively enriched, while iDOT1L-upregulated genes were positively enriched upon AF10 loss (**Figure 4B**). Several commonly regulated genes such as *EPCAM, COL6A2* and *NR2F2* were verified by qPCR (**Figure 4C**). Taken together, these data suggest that AF10 suppression and DOT1L inhibition have similar transcriptional effects during reprogramming. We functionally tested this notion by combining AF10 suppression with DOT1L inhibition. Individually, DOT1L inhibition or genetic suppression of AF10 increased reprogramming efficiency as expected; however, the combination of these perturbations did not result in a further increase in efficiency (**Figure 4D, E**). We also generated combined knock-out lines of both AF10 and DOT1L, verified the decrease in H3K79 methylation and then reprogrammed the resulting double knock-out cells (**Figure S1D**). AF10 and DOT1L double knockout did not significantly increase reprogramming compared to targeting each factor alone (**Figure 4F**). Overall, these results indicate that suppression of AF10 increases reprogramming mainly through its effect on DOT1L and H3K79 methylation.

**Figure 4:**
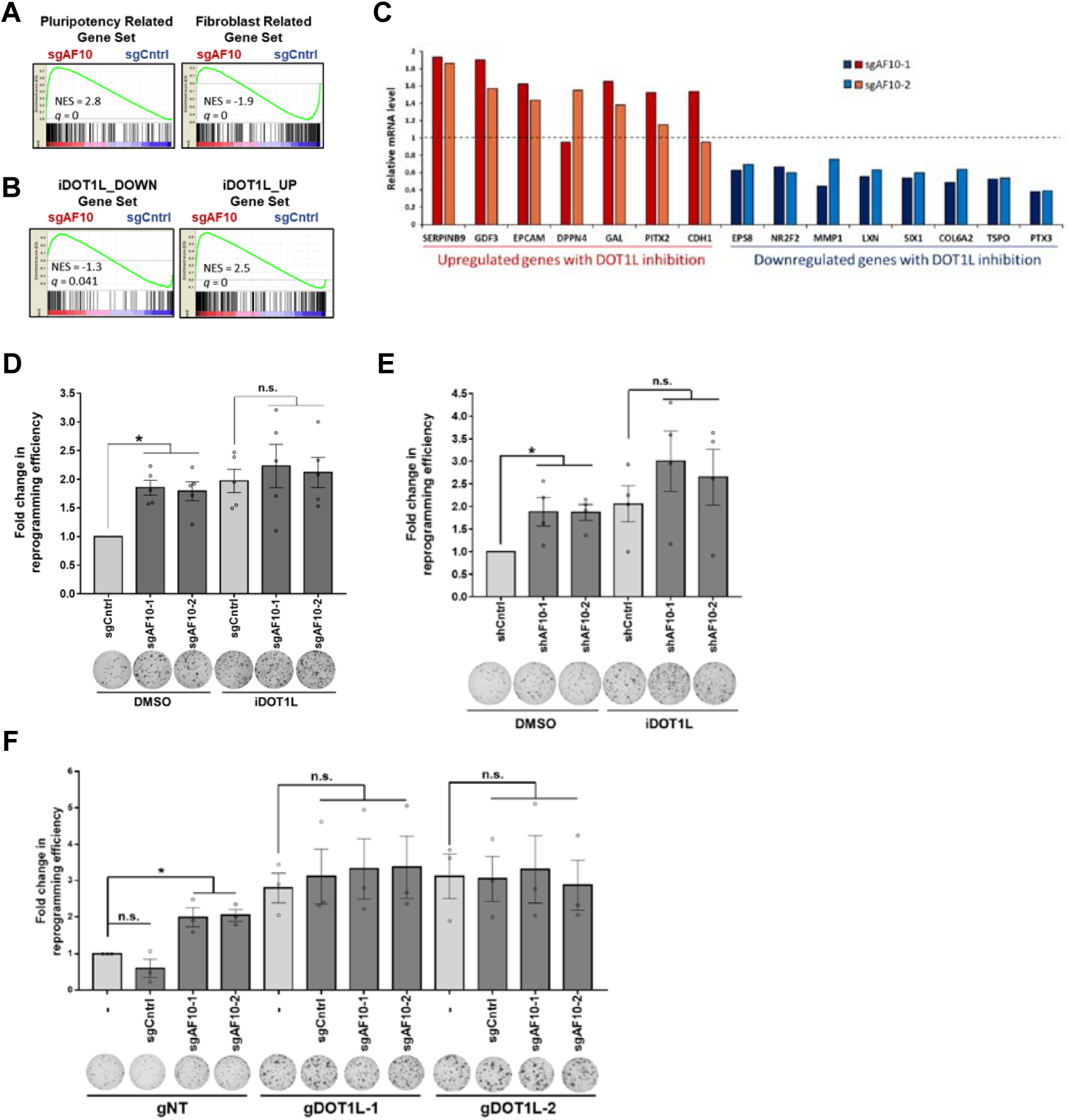
AF10 expression maintains somatic cell identity **(A)** Gene set enrichment analysis (GSEA) of transcriptome data of sgAF10 cells with respect to pluripotency-related and fibroblast-related gene sets. NES: normalized enrichment score, q val: False discovery rate (FDR) q-value. **(B)** Gene set enrichment analysis (GSEA) of transcriptome data of sgAF10 cells with respect to iDOT1L_DOWN and iDOT1L_UP gene sets. NES: normalized enrichment score, q val: False discovery rate (FDR) q-value. **(C)** mRNA levels for a set of DOT1L-regulated genes in AF10 knockout fibroblasts as determined by qRT-PCR. β-actin was used as an internal control and gene expression levels were normalized to sgControl expressing fibroblasts (dashed line). **(D)** Fold change in the number of Tra-1-60-positive colonies derived from AF10 sgRNA expressing cells after reprogramming in the presence of DMSO or a DOT1L inhibitor (iDOT1L; EPZ004777). *P* values were determined by one sample t-test; * *P*⍰< ⍰0.05. Bar graphs show the mean and error bars represent SEM in independent biological replicates (each circle). Representative Tra-1-60 stained wells are shown below the graph. *P* values were 0.001 for sgAF10-1 and 0.004 for sgAF10-2. n.s., not significant. **(E)** Fold change in the number of Tra-1-60-positive colonies derived from control or AF10 knockdown cells (shAF10) treated with either vehicle (DMSO) or iDOT1L (EPZ004777). Representative Tra-1-60-stained wells are shown below the graph. *P* values were determined by one sample t-test; * *P*1<10.05. Bar graphs show the mean and error bars represent SEM from independent biological replicates (each circle). *P* values were 0.034 for shAF10-1 and 0.007 for shAF10-2. n.s., not significant. **(F)** Fold change in the number of Tra-1-60-positive colonies derived from double knockout cells expressing DOT1L and AF10 targeting sgRNAs. *P* values were determined by one sample t-test; * *P*⍰< ⍰0.05. Bar graphs show the mean and error bars represent SEM in independent biological replicates (each circle). Representative Tra-1-60 stained wells are shown below the graph. *P* values were 0.031 for sgAF10-1 and 0.014 for sgAF10-2. n.s., not significant.

## Discussion

Here, we identified DOT1L-proximal proteins via proximity labelling and tested the effects of these proteins on somatic cell reprogramming. BioID-based proteomics uncovered TPR, KAISO, NUMA1, MRE11, NONO and SIN3B as novel DOT1L-proximal proteins in addition to known direct interactors of DOT1L, including AF10, AF17, ENL, Histone H1 and DDX21 (Mohan et al., 2010; Park et al., 2010; Wu et al., 2020). We tested the effect of DOT1L-proximal proteins in somatic cell reprogramming via loss of function experiments and showed that AF10 and NONO play functionally important roles in the generation of human iPSCs. Among these proteins, only loss of AF10 affected overall H3K79 methylation levels prompting us to further investigate its mechanism. In addition, several well-characterized DOT1L binding partners such as, AF17 and ENL had no effect in reprogramming pointing to a specific role for AF10 in this process, a finding corroborated in recent studies of mouse reprogramming (Wille et al., 2020). The fact that AF9, AF17 and ENL are members of Super Elongation Complex (SEC)(Wang et al., 2016), suggest that DOT1L’s role in suppressing cellular reprogramming is largely independent of its association with transcriptional elongation.

AF10 is a rate-limiting cofactor for higher order (di- and tri-) methylation of H3K79 and directly interacts with DOT1L through its octamer motif-leucine zipper (OM-LZ) domain (Chen et al., 2015b; Deshpande et al., 2014; Song et al., 2019). We show that this interaction is critical for AF10’s ability to prevent reprogramming. Furthermore, combined genetic suppression of AF10 and DOT1L did not result in an additive enhancement of reprogramming. Another potential function of AF10 is to act as a histone reader, recognizing unmethylated H3K27 and recruiting DOT1L to loci devoid of H3K27 modifications (Chen et al., 2015b). However, we find that histone-binding function of AF10 is not necessary to suppress reprogramming. Therefore, AF10 acts as a key barrier to reprogramming not through histone binding, but by regulating higher order H3K79 methylation by DOT1L.

AF10 suppression in somatic cells decreases results in wide-ranging dysregulation of gene expression during reprogramming. In particular, silencing of somatic-specific genes is facilitated by suppression of AF10, a finding in consonance with the effect of DOT1L inhibition. These findings indicate that AF10 acts as a safeguarding mechanism for somatic cell identity by enabling higher order H3K79 methylation of somatic-specific genes. Presence of higher order H3K79 methylation may antagonize gene repression, thereby preventing silencing of somatic transcriptional programs upon OSKM expression (Stulemeijer et al., 2011)(Aslam et al., 2021). Alternatively, recent work points to a role for DOT1L in transcription initiation, and it will be interesting to investigate if AF10 plays a role in that process (Wu et al., 2020). While the role of H3K79 methylation in preventing reprogramming to pluripotency is now well established, it will be of interest to test whether AF10 and DOT1L also regulate direct lineage conversions between terminally differentiated cells.

### Experimental Procedures

#### Plasmids

BirA^R118G^ (BirA*) cDNA was amplified from pcDNA3.1-mycBioID (Addgene, catalog no. 35700). DOT1L wildtype (WT) and mutant (G163R/S164C/G165R) cDNAs were described previously (Okada et al., 2005). In-frame BirA*-DOT1L fusion protein coding sequence was cloned into pENTR1A no ccDB (Addgene, catalog no. 17398) and transferred into expression plasmid pLEX-307 (Addgene, catalog no. 41392) via LR cloning (Invitrogen). pBabe-puro-AF10 wild-type (wt) and L107A mutant (mut) plasmids were gifts of Or Gozani (Stanford University). Wt- and mut-AF10 cDNAs were amplified with Phusion polymerase and inserted into pENTR1A no ccDB (Addgene, catalog no. 17398). OM-LZ domain (703-784) deleted cDNAs were prepared with Q5-site directed mutagenesis kit (NEB) according to manufacturer’s instructions. All AF10 sequences were cloned into a lentiviral expression plasmid pLenti CMV/TO Hygro DEST (Addgene, catalog no. 17291) via LR cloning (Invitrogen).

### shRNA and gRNA Cloning

shRNAs were designed and cloned into the MSCV-PM vector as previously described (Onder et al., 2012). All vectors were confirmed by Sanger sequencing. sgAF10 plasmids were gifts of Or Gozani (Stanford University). Rest of the gRNAs were designed and cloned into lentiCRISPRv2 (Addgene, catalog no. 52691) vector as previously described (Sanjana et al., 2014). shRNA and sgRNA sequences are listed in **Table S1**. All vectors were confirmed by Sanger sequencing using U6 promoter sequencing primer (5’-ACTATCATATGCTTACCGTAAC-3’).

### Reprogramming Assays

Fifty thousand dH1f cells (Park et al., 2008) were seeded onto 12-well plates and infected with lentiviral OSKM vectors (Addgene, catalog no. 21162, 21164). Medium was changed every other day with D10 medium (1XDMEM with 10% FBS, 1% Penicillin/Streptomycin). On day 6, cells were trypsinized and transferred onto mitomycin-c treated MEFs. Medium was then changed to hESC medium (DMEM/F12 with 20% KOSR, 1% L-glutamine, 1% non-essential amino acids, 0.055⍰mM beta-mercaptoethanol, 10⍰ng⍰ml^−1^ bFGF). Plates were fixed and stained for Tra-1-60 on day 21. iDOT1L (EPZ004777, Tocris) was used at 3⍰μM concentration for 6 days after OSKM infection.

### Production of Viral Supernatants

HEK-293T cells were plated at a density of 2.5⍰×⍰10^6^ cells per 10-cm dish and transfected with 2.5⍰µg viral vector, 2.25⍰µg pUMVC (Addgene, catalog no. 8449) for retroviruses or pCMV-dR8.2 ΔVPR (Addgene, catalog no. 8455) for lentiviruses with 0.25⍰µg pCMV-VSV-G (Addgene, catalog no. 8454) using 20⍰µl FUGENE 6 (Promega) in 400⍰µl DMEM per plate. Supernatants were collected 481hr and 721hr post-transfection and filtered through 0.45-µm pore size filters. To concentrate the viruses, viral supernatants were mixed with PEG8000 (Sigma, dissolved in DPBS, 10% final concentration) and left overnight at 4⍰°C. The next day, supernatants were centrifuged at 2500⍰rpm for 20⍰min, and pellets were re-suspended in PBS. Viral transductions were carried out overnight in the presence of 8⍰µg⍰ml^−1^ protamine sulfate (Sigma). Transduced cells were selected with 1⍰μg⍰ml^−1^ puromycin or 2001μg⍰ml^−1^ hygromycin.

### Generation of DOT1L-KO Single Cell Clones

HEK293T cells were transfected with either non-targeting (gCntrl) or guideDOT1L (gDOT1L) containing lenticrisprV2 plasmids and transfected cells were selected with 2 μg⍰ml^−1^ puromycin. After selection, cells were trypsinized, diluted to a single cell suspension and seeded onto 96-well plates. Single cell clones were identified and expanded. H3K79me2 levels in selected single cell clones were assayed via immunoblotting.

### Quantitative RT–PCR analyses

Total RNA was extracted using NucleoSpin RNA kit (Macherey Nagel) and reverse transcribed with Hexanucleotide Mix (Roche). The resulting complementary DNAs were used for PCR using SYBR-Green Master PCR mix (Roche) and run on a LightCycler 480 Instrument II (Roche) with 40 cycles of 10⍰s at 95⍰°C, 30⍰s at 60⍰°C and 30⍰s at 72⍰°C. All quantifications were normalized to an endogenous β-actin control. The relative quantification value for each target gene compared to the calibrator for that target is expressed as 2^−(Ct⍰−⍰Cc)^ (Ct and Cc are the mean threshold cycle differences after normalizing to β-actin). List of primers are in Table S1.

### RNA Sequencing and Analysis

RNA isolation was performed with Direct-zol kit (Zymo Research). NEBNext Poly(A) mRNA Magnetic Isolation Module from NEBNext Ultra Directional RNA Library Prep Kit for Illumina was used to enrich mRNA from RNA-sequencing samples. Samples were then validated on a Tapestation (Agilent) to determine library size and quantification prior to paired-end (2 × 41 bp) sequencing on a NextSeq 500 (Illumina) platform. Reads were mapped to hg19 built-in genome by HISAT2 after assessing their quality by FastQC. RNA-sequencing data are deposited to the NCBI GEO database with the accession number **GSE161043**. DeSeq2 package was used to find differentially expressed genes between samples. Genes were considered as differentially regulated based on |log2 fold change|> 0.5 and adjusted *p*-value<0.05. Differential gene expressions between pluripotent stem cells and fibroblast cells were computed by affy and limma packages from R to generate fibroblast- and pluripotency-related gene sets as described previously (Ebrahimi 2019). Differential gene expression analysis to generate iDOT1L regulated gene sets is performed on GEO2R web tool between dH1f-inhibitor-OSKM samples and dH1f-untreated-OSKM samples from GSE29253 (Onder et al., 2012). iDOT1L_UP gene set is composed of genes that are upregulated in treatment group (*p*-value<0.05 and logFC>0.5) and iDOT1L_DOWN gene set is composed of genes that are downregulated in treatment group (*p*-value<0.05 and logFC<-0.5). Rank-ordered gene lists were used for gene-set enrichment analysis (Subramanian et al., 2005).

### Nuclear Protein Extraction and Histone Acid Extraction

Cell pellets were resuspended in cytosolic lysis buffer (10 mM HEPES pH7.9, 10 mM KCl, 0.1 mM EDTA, 0.4% NP-40, cOmplete ULTRA protease inhibitor Tablets [Roche]) and incubated for 15 min on ice and centrifuged at 4°C for 3 min at 3000*g*. Pellets were washed once with cytosolic lysis buffer and then resuspended in nuclear lysis buffer (20 mM HEPES pH7.9, 0.4 M NaCl, 1 mM EDTA, 10% Glycerol, cOmplete ULTRA protease inhibitor Tablets [Roche]) followed by sonication 2 times for 10 seconds at 40 amplitude with a 10 second interval in between (QSONICA Q700 with microtip). After sonication, tubes were centrifuged at 4°C for 5 min at 15000*g*. Supernatant was removed as nuclear protein fraction. For histone acid extraction, cell pellets were resuspended with triton extraction buffer (0.5% Triton X-100, 2mM PMSF, 0.02% NaN_3_ in PBS) and incubated for 10 min on ice then centrifuged at 4°C for 10 min at 2000rpm. Pellet was washed with triton extraction buffer and centrifuged again. Supernatant was discarded and the pellet was resuspended in 0.2N HCl. Tubes were incubated at 4°C for 16 hours on a rotating wheel and centrifuged at 4°C for 10 min at 2000rpm. Supernatants were neutralized with the addition 0.1M NaOH for 1/5 volume of HCl solution. Protein concentrations were determined via BCA assay (Thermo Scientific).

### Immunoblotting

Equal amounts of proteins were boiled with loading buffer (4X Laemmli sample buffer, Bio-Rad) and loaded onto 4–15% Mini-PROTEAN TGX Precast Protein Gels (Bio-Rad). Gels were run with TGS buffer (diluted from 10X stock, Bio-Rad). Precision Plus Protein Dual Color Standards (Bio-Rad) were used a molecular weight ladder. Proteins were transferred onto Immun-Blot PVDF Membrane (Bio-Rad) via Trans-Blot Turbo Transfer System (Bio-Rad). Membrane was incubated with 5% blotting grade blocker (Bio-Rad) dissolved in TBS-T (20 mM Tris, 150 mM NaCl, 0.1% Tween 20 –pH 7.6). For Streptavidin-HRP blotting membranes were blocked with 2% bovine serum albumin (BSA, Sigma) in TBS-T. Primary antibodies were incubated on membranes at 4°C for 16 hours. Primary antibodies were Streptavidin-HRP (BioLegend 405210, 1:10,000), H3K79me2 (ab3594, 1:1000), H3 total (ab1791, 1:1000). After primary antibody incubation, membranes were washed and then incubated with secondary antibody solution (1:5000 secondary antibody ab97051 in 5% blotting grade blocker in TBS-T) at room temperature for 1-2 hours. Membranes were washed with TBS-T and proteins were visualized with Pierce ECL Western Blotting Substrate (Thermo Scientific) and Odyssey Fc Imaging systems (LiCor).

### Pull-Down assays and Mass Spectrometry Analysis for BioID

HEK-293T cells were infected with lentiviral BirA*-DOT1L. Puromycin selected cells were expanded and incubated with 50 μM D-Biotin (Sigma, 47868) for 24 hours. Proteins were obtained via nuclear fractionation method. As a control, uninfected HEK293T cells were used. Pull-down was performed with Streptavidin beads (Thermo Scientific, 53117) as previously described (Firat-Karalar et al., 2014). Briefly, 3 mg nuclear fraction was incubated with 100 μl Streptavidin beads at 4°C for 16 hours on a rotating wheel at 10 rpm. Then supernatants were collected, and beads were washed twice in 2% SDS; once with wash buffer 1 (0.2% deoxycholate, 1% Triton X, 500 mM NaCI, 1 mM EDTA, 50 mM HEPES, pH 7.5), once with wash buffer 2 (250 mM LiCI, 0.5% NP-40, 0.5% deoxycholate, 1% Triton X, 500 mM NaCI, 1 mM EDTA, 10 mM Tris, pH 8.1) and twice with wash buffer 3 (50 mM Tris, pH 7.4, and 50 mM NaCI). Eluted proteins were analyzed with Streptavidin-HRP antibodies to observe the efficiency of pull-down. For mass spectrometry analysis, control (uninfected) and BirA*-AF10 WT or MUT expressing HEK293T cells were used. Following nuclear protein isolation and streptavidin pulldown, bound proteins were digested with on-bead tryptic proteolysis as previously described (Özkan Küçük et al., 2018). Briefly, beads were washed (8 M urea in 0.1 M Tris-HCl, pH 8.5) and reduction and alkylation steps performed. After a final wash with 50 mM ammonium bicarbonate, beads were treated with trypsin overnight. Reaction was quenched with acidification and the resulting peptides were desalted (Rappsilber et al., 2003) and then analyzed with reversed-phase nLC (NanoLC-II, Thermo Scientific) combined with orbitrap mass spectrometer (Q Exactive Orbitrap, Thermo Scientific). The raw files were processed with Proteome Discoverer 1.4 (Thermo Scientific) using human Uniprot database (Release 2015-21,039 entries) as previously described (Kagiali et al., 2019; Özkan Küçük et al., 2018). Two technical replicates were performed for each sample. To identify DOT1L-specific biotinylation, proteins detected in HEK293T control samples were subtracted from BirA* infected samples. The remaining proteins were selected only if were present in both runs of mass-spectrometry. Among these common proteins, nuclear localized ones are determined via GO annotation (http://www.geneontology.org/) using cellular component analysis. UniProt protein names were converted via ID mapping tool (https://www.uniprot.org/uploadlists/). Determined proteins were sorted according to their sequence coverage and abundance using PSM (peptide spectrum matches) numbers.

### Tra-1-60 Staining and Quantification

To quantify the number of iPSC colonies, reprogramming plates were stained with Tra-1-60 antibody as previously described (Onder et al., 2012). Briefly, cells were fixed with 4% paraformaldehyde and incubated with biotin-anti-Tra-1-60 (BioLegend, catalog no. 330604, 1:250) diluted in PBS with 3% FBS and 0.3% Triton X-100. Followed by incubation with streptavidin-HRP (Biolegend, catalog no. 405210, 1:500). Staining was developed with the DAB peroxidase substrate solution (0.05% 3,3’-diaminobenzidine [Sigma, D8001], 0.05% nickel ammonium sulfate and 0.015% H^2^O^2^ in PBS, pH 7.2) and iPSC colonies were quantified with ImageJ software (https://imagej.nih.gov/ij/).

### T7-endonuclease assay

gRNA infected cells were harvested, and genomic DNAs were isolated using MN Nucleospin Tissue kit. gRNA targeting sites were amplified with specific primers (Table S1) PCR clean-up was performed (MN, PCR clean up and gel extraction kit). 400 ng from cleaned PCR products were mixed with NEB 2 buffer and incubated according to heteroduplex formation protocol (5 minutes at 95°C and ramp down to 85°C at -2°C/sec and ramp down to 25°C at -0.1°C/sec). After heteroduplex formation, samples were treated with T7 endonuclease (NEB) for 1-2 hours at 37°C. Digested samples were analyzed on 2% agarose gels and visualized via Gel Doc XR System (Bio-Rad).

### Teratoma formation assay

All experiments were carried out under a protocol approved by Koç University Animal Experiments Ethics Committee. Injections were performed as previously described (Fidan et al., 2015). Briefly, iPSCs from 80% confluent 10 cm dish were collected using ReLeSR (Stemcell Technologies) and re-suspended in 100 μl ice-cold 1:1 mixture of Matrigel (Corning) and hES growth medium. Intramuscular injections were performed in SCID mice. Teratomas were collected 8–10 weeks after injection and analyzed histologically via hematoxylin and eosin staining.

### Immunofluorescence staining

Immunostainings were performed as previously described (Ebrahimi et al., 2019). Briefly, iPSCs from single cell clones were fixed with 4% paraformaldehyde in PBS and incubated overnight at 4°C with primary antibody: OCT4, (Abcam, ab19857), SSEA4 (BD, 560219), NANOG (Abcam, ab21624). Nuclei were stained with DAPI (Vectashield, H-1500). Images were acquired using a Nikon 90i confocal microscope.

## Acknowledgments

We would like to thank Ahmet Kocabay and Ali Cihan Taşkın for help with mouse experiments. The authors gratefully acknowledge use of the services and facilities of the Koç University Research Center for Translational Medicine (KUTTAM), funded by the Republic of Turkey Ministry of Development. The content is solely the responsibility of the authors and does not necessarily represent the official views of the Ministry of Development. This work was supported by EMBO Installation Grant (TO), Newton Advanced Fellowship (T.O.), TUBITAK Project <u>115Z706</u> (T.O.), Arthritis Research UK (program grant 20522, UO), Cancer Research UK (UO) and the LEAN project of the Leducq Foundation (UO). The research leading to these results has received funding from the People Programme (Marie Curie Actions) of the European Union’s Seventh Framework Programme (FP7/2007-2013) under REA grant agreement n° [609305].

## Author Contributions

Conceptualization, D.U.Ç. and T.T.O,; Methodology D.U.C, N.E.Ö.K,; Investigation D.U.Ç., D.O., B.Ö., C.A., M.P., D.D.; Formal Analysis D.U.Ç., K.S.; Resources, U.O., and N.Ö; Supervision, T.T.O

## Declaration of Interests

The authors declare no competing interests

## Supplemental Information

**Supplementary Figure 1:** Validation of AF10 inhibition in somatic cells and iPSCs

**(A)** mRNA levels of shRNA targeted genes were assessed via qRT-PCR. β-actin was used as an internal control and gene expression levels are normalized to control shFF (firefly luciferase targeting shRNA) expressing cells.

**(B)** Immunoblot for H3K79me2 in shRNA-targeted fibroblasts. Total H3 levels were used as loading control.

**(C)** AF10 mRNA levels in individual iPSC clones derived from control and AF10 sgRNA expressing fibroblasts as determined by qRT-PCR. β-actin was used as an internal control and expression level is normalized to sgControl-1 iPSCs.

**(D)** Immunoblot for H3K79me2 levels in double sgRNA expressing fibroblasts. Total H3 levels were used as loading control. gNT: non-targeting gRNA; gD1/2, DOT1L targeting gRNA 1 or 2.

**Supplementary Table 1**. List of oligonucleotides for cloning and PCR

**Supplementary Table 2**. Raw data of mass spectrometry analysis of BioID assay

## Notes

### Competing Interest Statement

The authors have declared no competing interest.

### Summary of Updates

Discussion section added

