## Supplementary figures and images for "AF10 (MLLT10) prevents somatic cell reprogramming through regulation of DOT1L-mediated H3K79 methylation"

### Supplementary Figures

#### Slide 1
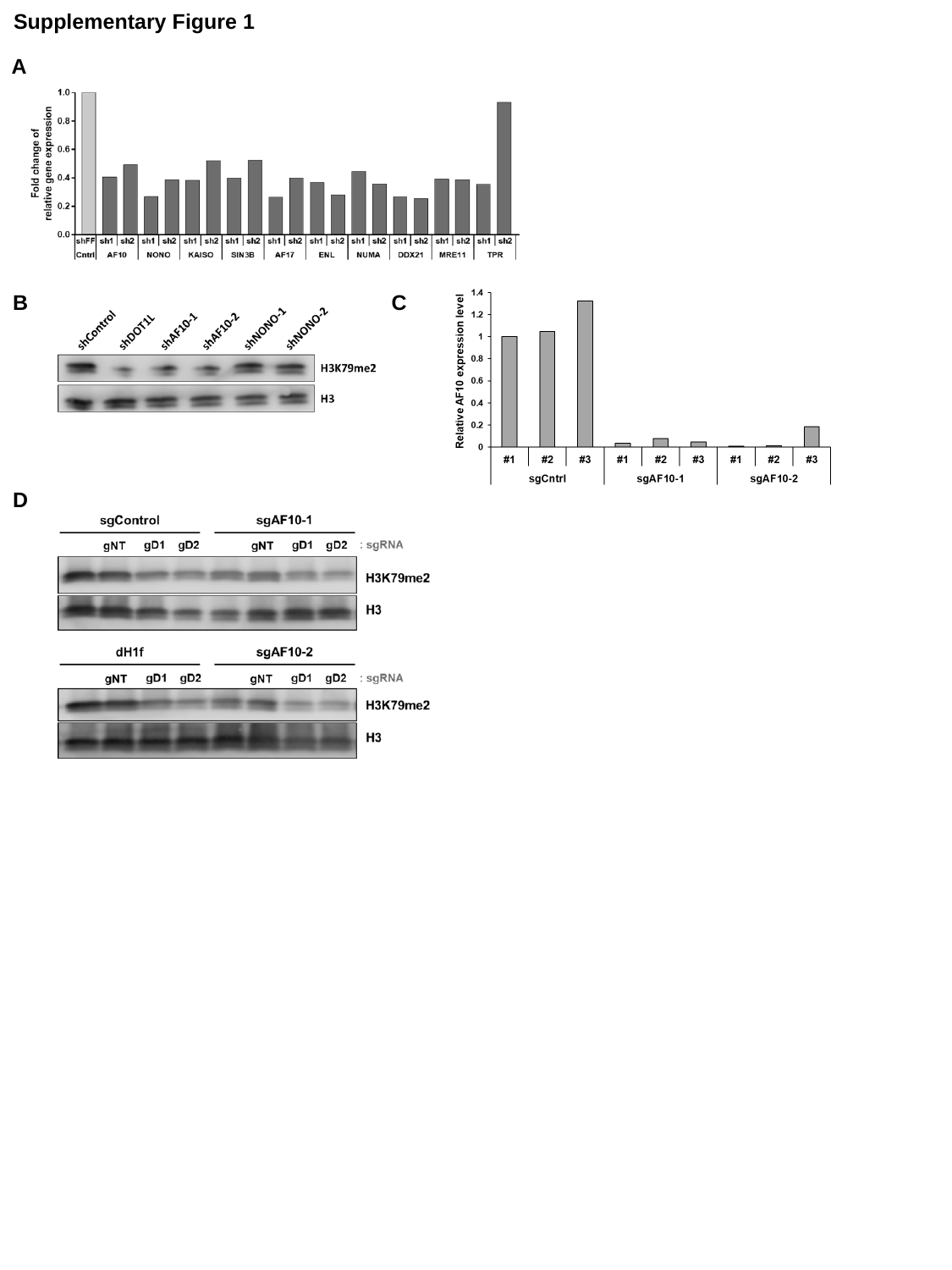

Supplementary Figure 1
A
B
C
D
